# Transient structural properties of the Rho GDP-dissociation inhibitor

**DOI:** 10.1101/2023.09.13.557459

**Authors:** Sara Medina Gomez, Ilaria Visco, Felipe Merino, Peter Bieling, Rasmus Linser

**Affiliations:** Department of Chemistry and Chemical Biology, TU Dortmund University, Otto-Hahn-Str. 4a, 44227 Dortmund, Germany; Department of Systemic Cell Biology, Max Planck Institute of Molecular Physiology, Dortmund; Department of Protein Evolution, Max Planck Institute of Developmental Biology, Tübingen

## Abstract

Rho GTPases are master spatial regulators of the cytoskeleton that control a wide range of cellular processes. Their inactivation by removal from cellular membranes involves the stepwise formation of a stable complex with guanine nucleotide dissociation inhibitors (RhoGDIs), for which process the RhoGDI N-terminus is indispensable. The formation of this interface has been thought to emerge from an intrinsically disordered state of RhoGDI in its free, apo form. Here we use tailored solution NMR analyses, molecular dynamics simulations, and biochemical essays to pinpoint the site-specific structural features of full-length RhoGDI1 before and after binding its GTPase client Cdc42. In contrast to the current mechanistic understanding, a diverse set of NMR data unequivocally shows that the structural properties of the GDI N-terminus seen in crystal structures of the complex with GTPases already exist as largely preformed features in free, apo GDI. Even more interestingly, the required structural properties are imposed onto the terminus context-specifically by modulating interactions with the surface of the folded C-terminal domain. Lastly, upon Cdc42 binding, the flexibility of the N-terminus and its secondary-structural propensities are not largely abrogated. These observations change the textbook picture of the mechanism of membrane extraction of the GTPase. Rather than a disorder-to-order transition upon binding, an active role of the N-terminus with differentially preformed structural properties, suitably modulated by the specific surrounding along the multi-step process, seems required to leverage the intricate and highly selective extraction process.

## Introduction

Small GTPases of the Ras superfamily are key membrane-associated signaling molecules that can assume distinct activity states, which depends on the regulated hydrolysis and exchange of their associated guanine nucleotide. The Rho family of small GTPases in particular are responsible for locally modulating cytoskeleton dynamics in intracellular organization, cell polarity, morphogenesis, motility and other essential cellular processes.^1, 2^ Rho guanine nucleotide dissociation inhibitors negatively regulate RhoGTPases by extracting them from membranes^3^, sequestering the membrane-binding carboxy-terminal isoprene moiety^4, 5^ and suppressing interactions with regulators of the GTPase activity states such as nucleotide exchange factors (GEFs) or GTPase-activating proteins (GAPs, see Figs. 1A and B).^4, 5, 6^ Thereby, RhoGDIs maintain a large pool of soluble, inactive RhoGTPases. The highly conserved N-terminus of RhoGDI (Fig. 1C) is known to be necessary for membrane extraction^3^ but has also been thought of as responsible for inhibition of both nucleotide exchange and hydrolysis by sterically blocking the switch regions of its GTPase clients, thereby locking them in an inactive state.^4, 6, 7, 8^ It also contributes significantly to the binding energy of the RhoGDI:RhoGTPase complex, since its complete removal reduces affinity for GTPases by more than 100-fold.^6^ Its functional importance is also in line with the high level of sequence conservation between distinct RhoGDI orthologs and isoforms (Fig. 1C). The N-terminus in the apo protein is, however, thought of as disordered,^5, 6, 9^ as it has low binding affinity on its own, no significant NOEs were observed, and the N-terminal resonances cluster in the central region of the NMR HSQC spectra, a behavior typical for intrinsically disordered protein regions lacking defined structural features. Fig. 1D shows the prediction of order/disorder purely on the basis of in-silico assessment using the ODINpred server.^10^

**Fig. 1:**
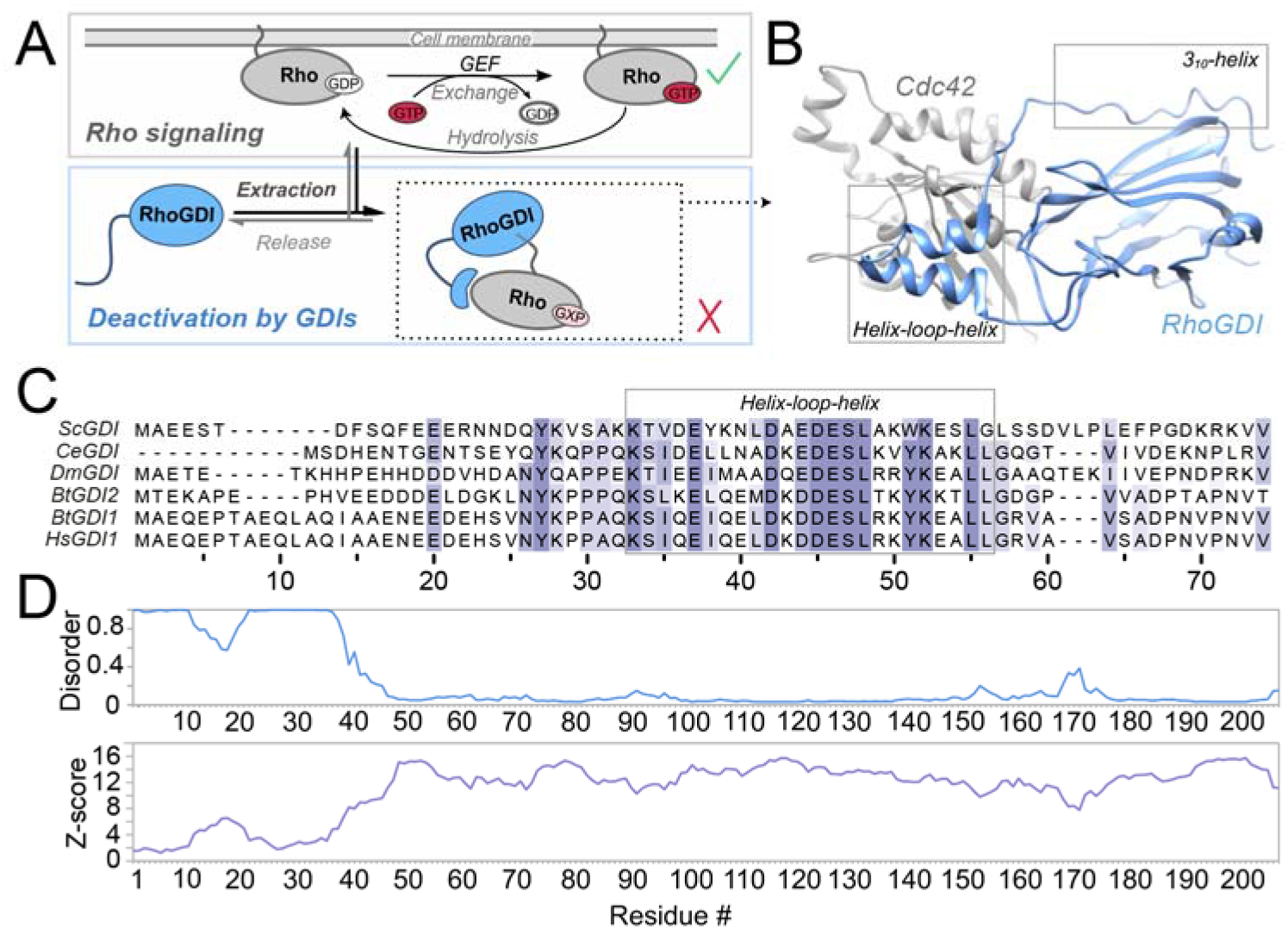
RhoGDI function and bioinformatical analysis. **A**) RhoGDI deactivates RhoGTPases by their membrane extraction, upon which a high-affinity complex is formed. “GXP” denotes that both, active (GTP-bound) and inactive (GDP-bound) Rho can be extracted.^3^ **B**) Crystal structure of RhoGDI in complex with the GTPase Cdc42 (PDB 1DOA), showing the specific tertiary and secondary-structural features newly adopted by the previously flexible N-terminus (gray boxes). **C**) Sequence alignment of the N-terminal region of different RhoGDI orthologs and isoforms (also compare Fig. S1). Sc: *Saccharomyces cerevisiae*, Ce: *Caenorhabditis elegans*, Dm: *Drosophila melanogaster*, Bt: *Bos taurus*, Hs: *Homo sapiens*. The color depicts the BLOSUM62 score. Val74 marks the start of the C-terminal Ig-like domain. **D**) Disorder prediction (blue) and secondary-structure Z-score (purple) from pure *in silico* assessment using ODiNPred,^10^ demonstrating the dominantly disordered character of the N-terminal primary sequence in consistency with the original literature.^5, 6, 9^

Upon complex formation with GTPases, the GDI N-terminus, by contrast, is thought to fold into a structured configuration in an induced-fit manner, now becoming stably bound to both the GTPase and parts of its own folded protein core,^4, 7, 8^ see the corresponding crystal structure Fig. 1B. In particular, the folded conformation now contains a helix-loop-helix motif, which contacts switch I and II regions of the RhoGTPase, and a partially helical structure for the very N-terminus, including a 3^10^-helix for residues 9 to 16, which folds back against the immunoglobulin (Ig)-like domain that contributes residues to the geranylgeranyl-binding pocket.^4^ This conversion of the secondary structure of the N-terminus from disorder to order is thought to be an integral part of the difficile mechanism to extract RhoGTPases from the membrane. Based on existing structural^4, 7, 8^ and biochemical data, a two-step mechanism has been proposed in which the unstructured GDI N-terminus first folds onto the membrane-bound GTPase.^11^ A subsequent isomerization event then leads to the swapping of the prenyl moiety between the membrane and the Ig-like domain of RhoGDI. How the extreme extent of alteration in the structural properties of the RhoGDI N-terminus can be reconciled, however, with its role in the rapid and specific recognition of membrane-bound GTPases is presently unclear. Moreover, its character as a disordered peptide in apo GDI, its low binding affinity to the GTPase in isolated form, and the well-defined structural properties in the complex structure, where it folds into a location distant from the nucleotide, seem to contradict each other. Here we use NMR spectroscopy and biochemical assays to revise the understanding of structural properties for the N-terminal domain of RhoGDI in apo form and when bound to its RhoGTPase client Cdc42. Given its more widespread expression and higher affinity for its GTPase clients compared to other RhoGDI isoforms,^5, 12^ we restrict our analysis to RhoGDI1.

## Results

Our interest in the structural properties of the RhoGDI1 N-terminus emerged from an analysis via the current Robetta modeling routine, a deep-learning-based protein structure prediction tool.^13^ Even though such prediction results can potentially derive from other than apo states, it sparked our curiosity that the Robetta prediction for the N-terminus of the *apo* protein showed the presences of helical elements for residues 8 to 16 and the helix-loop-helix (Fig. 2A), in conflict with the current mechanistic model. (Fig. S2 compares predictions for the full-length protein with those for an isolated N-terminus.) To test *in silico* whether/how (partially) folded states are indeed possible within the conformational ensemble of RhoGDI’s N-terminal end and how this could be reconciled with the previous data, we first turned to molecular dynamics (MD) simulations with structure-based potentials. These simplified simulations are known to capture a protein’s folding process without the typical limitations imposed by conventional all-atom MD runs. As reference for the native state, we used the structure of RhoGDI observed in the complex with CDC42 (PDBID: 1DOA).

**Fig. 2:**
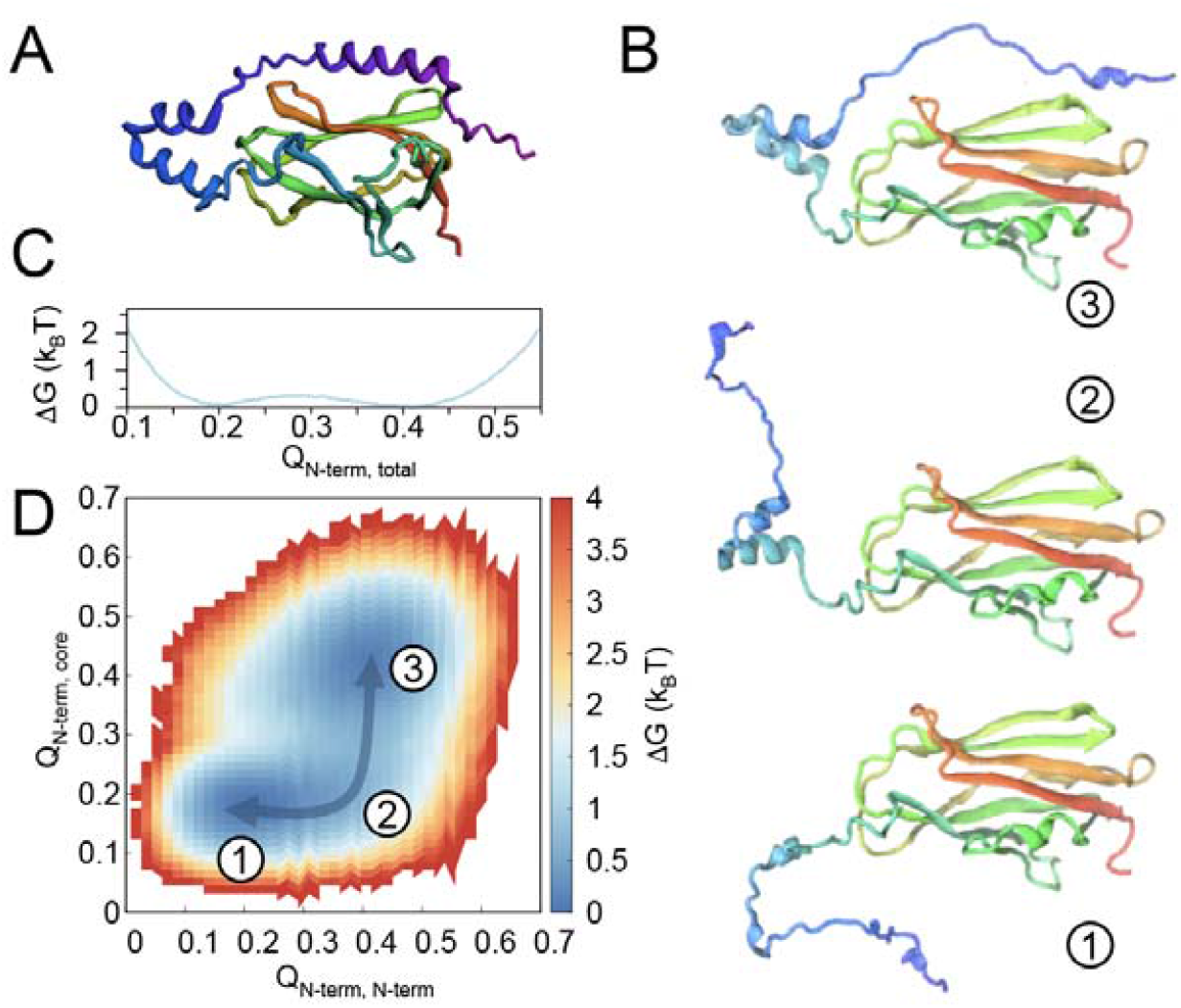
Structural properties from evolutionary analysis and in MD simulations. **A)** Robetta output, predicting strong N-terminal secondary-structural properties instead of the assumed disordered character. **B)** Representative snapshots of conformations visited by RhoGDI during the simulations. (Color in A) and B) according to primary sequence.) C and D) Folding free-energy profiles for RhoGDI”s N-terminus at its folding temperature: **C)** Free energy as a function of the fraction of all native contacts formed by the N-terminus, i. e., amino acids 1-65 (Q_Nterm, total_), and **D)** 2D free-energy profile of N-terminal folding as a function of the fraction of native contacts among N-terminal atoms (x-axis, Q_N-term, N-term_) and those between N-terminal atoms and the rest of the protein (y-axis, Q_Nterm, core_). The arrow highlights the minimum-free-energy path separating the fully folded from fully unfolded N-terminus.

Fig. 2B shows three representative conformations visited by the RhoGDI’s N-terminus during the simulations. Consistent with its loose connection to the rest of the protein, the N-terminus explores a large conformational ensemble, while the protein’s core domain remains folded. Folded and unfolded states are separated by a very shallow barrier, well below thermal energy (Fig. 2C). Interestingly, we consistently observed conversion between completely unfolded conformations and states where N-terminal secondary-structural elements (in particular the helix-loop-helix motif and the N-terminal 3_10_ helix) are spontaneously formed without attachment to the protein’s core. Indeed, the 2D folding free-energy profile as a function of the intra-N-terminal contacts and those between any N-terminal residues and the core shows that, while the expected free-energy minima representing the fully disordered and the fully attached state are present (states “1” and “3”, respectively, in Figs. 2B and D), a shallow path – the minimum free-energy path – connects them through an intermediate with partly folded, but detached N-terminus (state “2”).

To quantify the tendency towards a preformed N-terminal binding interface for GDI/GTPase interactions experimentally, we turned to NMR spectroscopic characterization of residue-specific residual structural and dynamics properties.^14, 15, 16, 17, 18, 19^ For that purpose, bovine RhoGDI1 was expressed and purified recombinantly (see details on expression and purification in the Materials and Methods) both in full length (fl, residues 1 to 204) and as an isolated N-terminal fragment (residues 1 to 59). To enable site-specific characterization of the secondary structural features, a comprehensive suite of 3D solution NMR spectra of the different RhoGDI1 constructs expressed in either doubly (^13^C, ^15^N) or triply labeled (^2^H, ^13^C, ^15^N) fashion were recorded. (See details in the Experimental Section.) As the N-terminal residues were known to cluster in a heavily overlapped central region of the H/N plane, compromising the assignment of the core residues in previous studies,^6^ we first turned to assignment of these residues in an isolated N-terminal fragment. Complete assignment of the N-terminal residues, their successive transfer to the full-length protein sample, and comprehensive assignment of the remaining residues in the full-length construct thus turned out to be possible. Figs. 3A and 4A show the assigned HSQC spectra for the entire RhoGDI1 protein and the isolated N-terminus, respectively. Chemical shifts were deposited into the BRMB under accession number 51835.

**Fig. 3:**
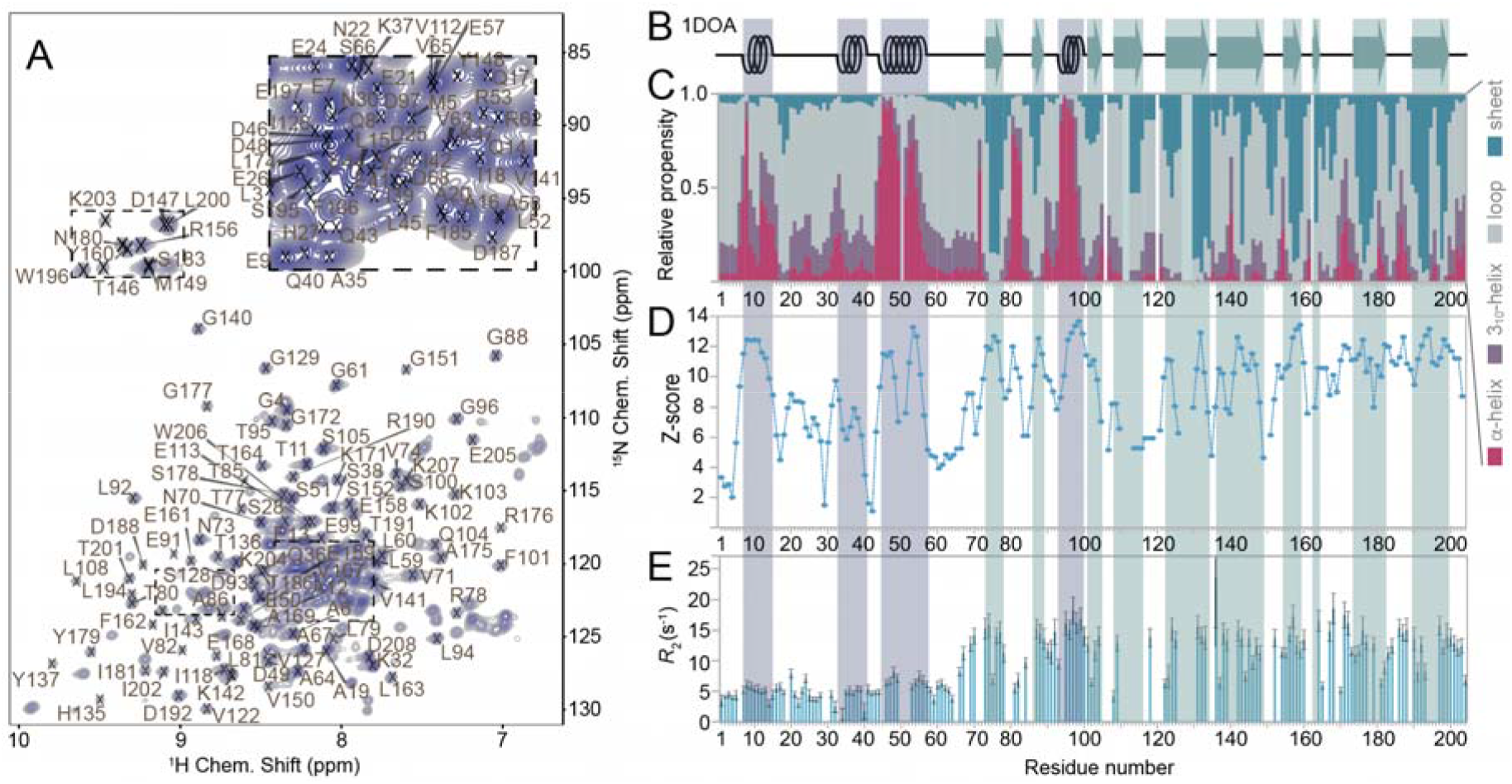
Experimental NMR-chemical-shift-based assessment of fl-RhoGDI1 secondary-structural propensity. **A**) Assigned HSQC of the full-length protein recorded at 800 MHz H Larmor frequency, with parts of the crowded central region only annotated in the magnified excerpts on top (dashed rectangles). **B**) Secondary structure found in the X-ray structure of the RhoGDI1:Cdc42 complex (PDB 1DOA). **C)** Chemical-shift-based relative secondary-structural propensities and **D**) Z-score of apo RhoGDI1 (from CheSPI). **E**) *R*_2_ relaxation of apo GDI, which confirms the existence of local (temporary) secondary structure in the sections denoted by CheSPI. The color code for secondary-structural analysis is shown on the right.

The chemical-shift assignments provided a springboard for experimental interrogation of RhoGDI1 structural features under close-to-physiological conditions in a site-specific manner. We first used the computational framework CheSPI (Chemical shift Secondary structure Population Inference^20^) for secondary-structural assessment based on actual backbone ^1^H^N, 15^N, ^13^CO, ^13^Cα and ^13^Cβ chemical shifts, which relies on neighbor-corrected chemical shifts and can distinguish between eight DSSP^21^ classes of temporarily adopted secondary structure. Opposed to secondary-structure prediction by TALOS^22^, regularly used for assessment of secondary structure in folded proteins, the accuracy of CheSPI prediction for secondary structure propensities provides a minimal bias with respect to folded structural elements and quantifies relative structural features in both ordered and disordered systems. (The outputs from Talos+ and ChesPI are specifically compared in Figs. S3 and S5.) The Z-score obtained from CheSPI analysis indicates the level of order/disorder as a function of sequence. A low Z-score value indicates a disordered region, while a high value indicates the presence of more stably adopted secondary structure. For an intrinsically disordered N-terminus, a low Z-score would be expected. Instead, the obtained result confirms the presence of pronounced secondary structure within the N-terminus of the apo form of the full-length protein (Fig. 3B-D). In particular, residues 42 to 57 display a stably helical fragment, exactly matching the helical structure found for the RhoGDI:GTPase complex.^4, 7, 8^ The second, shorter helical stretch seen in the complex, around residues 32 to 40, again shows helical secondary structural propensity, however, with much weaker helical (around 30 % overall) propensity and a lower Z-score (Fig. 3D) than the first one. A third helical fragment is found experimentally for residues 8-15 (the 3_10_-region in the complex), again with a quantitatively lower prediction propensity (around 50 %), but with a high Z-score. All of the (transient) helical propensities in the terminus align with the structural features of RhoGDI1 in the complex (Fig. 3B). Fig. 3E shows site-specific *R*_2_ rates of the apo-RhoGDI1. These rates (around 5 s^−1^, opposed to rates of around 15 s^−1^ for the structured residues in the core) confirm a greater overall flexibility (detachment and an individual tumbling correlation time) of the N-terminus, however, with slight but unequivocal elevations (7-8 s^−1^) for the transient secondary structure in the very N-terminus (around residue 8) and in particular for the more durable helix of the helix-loop-helix motif (around residue 50).

To decipher whether/to what extent the residual secondary-structural propensities found for the N-terminus are inherent to the primary sequence or rather determined by close proximity to interaction surfaces exposed by the protein core, assessment of secondary-structural features was performed for the isolated N-terminus (Fig. 4) as an unphysiological reference case. Whereas ^3^*J* HNHA couplings did not yield any consistent patterns (data not shown), for the very N-terminus, the CheSPI results (Fig. 4C) still qualitatively agree with the propensities of the full-length protein. (Disorder prediction from pure *in silico* assessment using ODiNPred^10, 20^ is shown for comparison in Fig. S4.) Also, the helical stretch seen for residues 8-15 has similarly low propensities, and the low helical propensity around residue 35 appears similar. The quantitative extent of helical propensity of the C-terminal residues of the helix-loop-helix motif (residues 45 – 55), however, strongly differs, with an only minor degree of helicity in the isolated terminus (around ∼10 % compared to > 90 % in the fl protein). To address potential imperfections of the chemical-shift-derived secondary-structural prediction for very low populations, the residual secondary-structural propensities witnessed were additionally probed by inspecting NOEs among and in between amide and Hα protons (Fig. 4D/E), which validate the transient formation of helical structure where expected. Opposed to the features of residues 8 – 15, the differences between isolated N-terminus and full-length protein for the long helical stretch of the final binding interface to the GTPase suggest that it is the (transient) interaction with the core that triggers the rather stable preformation of desired secondary structural propensities, strongly aiding in prestructuring the GDI interface for the initiation of complex formation: Whereas on its own, the C-terminal part of the N-terminus (residues 30 – 60) is dominated by the features of an intrinsically disordered protein and shows only little tendency to form the final interface, in line with the previously observed inability of the isolated N-terminus to capture the client,^6^ the interactions of the N-terminus with the core – albeit not firmly attached – increase the population of transient secondary structural features to such extent that the interface for GTPase binding becomes preformed in a substantial manner. The comparison of chemical shifts between the isolated N-terminus and the full-length protein confirms these changes in the secondary structure (Fig. 4F). In addition to the residues in the helix-loop-helix region, experiencing large shifts, other residues with slight perturbations exist. Even though we restrict this analysis to the backbone due to the difficulty of sidechain assignments in the full-length protein, transient interactions with the core become obvious throughout the N-terminus, together strongly modulating the temporary structural features of the overall mobile terminus.

**Fig. 4:**
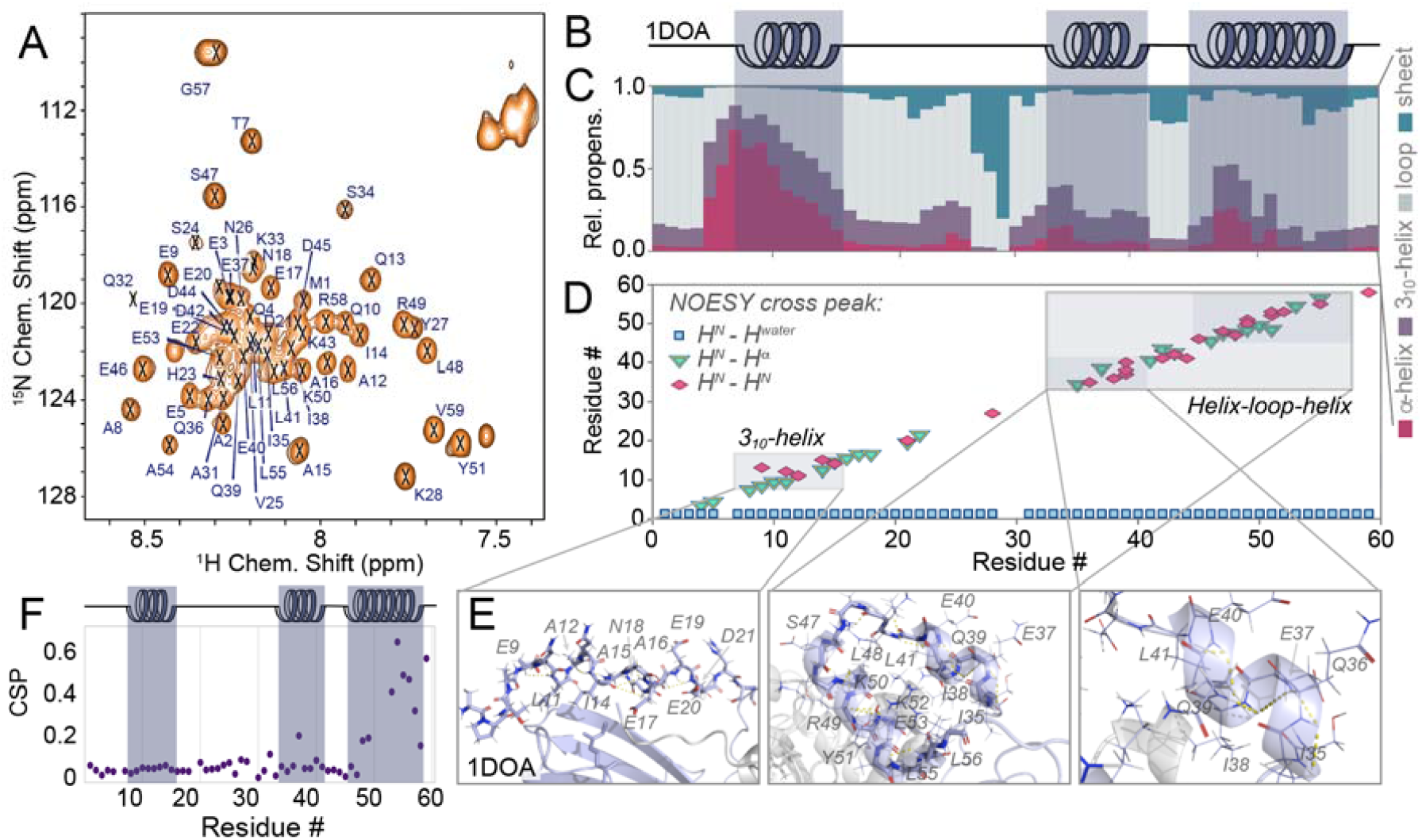
Characterization of the isolated RhoGDI1 N-terminus. **A**) Assigned ^1^H-^15^N HSQC of the isolated N-terminus (residues 1 to 59). **B**) Secondary structure of the X-ray structure (1DOA) of the complex. **C**) Analysis of the extent of secondary structure of the isolated RhoGDI N-terminus from backbone chemical shifts^20^. Color code depicted on the right. **D**) Presence of amide NOE cross peaks in a 3D ^15^N-edited NOESY experiment, showing the presence of transient secondary structure via magnetization transfer to other nearby amides (magenta diamonds), close-by H^α^ spins (cyan triangles), and to water (blue squares in the bottom). **E**) Depiction of matching secondary structure within the stretches of temporary secondary structure as seen in the complex with H-bonds depicted by yellow dashes. **F**) Chemical-shift perturbations between the isolated N-terminus and the full-length protein as a function of residue.

We next compared the transient structural properties that we observed in the apo protein with those of RhoGDI1 in complex with its binding partner GTPase CDC42 (Fig. 5). For this purpose, RhoGDI was quantitatively incorporated into a stoichiometric 1:1 complex by addition of excess Cdc42, which had been homogenously geranylgeranlyated *in vitro*. Given the sub-nM affinity of the complex (see below), the heterodimeric complex was separated from excess free Cdc42 by size exclusion chromatography. (See Materials and Methods for details of these procedures.) In order to verify complex formation, the *R*_2_ rates from the two samples were compared (Fig. 5C). *R*_2_ rates are modulated by the time scale of molecular tumbling (τ_c_), and an overall increase is expected when the effective molecular weight and thus τ_c_ increase. This tendency is clearly observed both, for the core residues (from 11.9 ± 3.9 to 19.6 ± 5.8) as well as for the N-terminus (from 5.2 ± 2.0 to 7.7 ± 3.1) when overlaying the *R*_2_ rates from both samples.

**Fig. 5:**
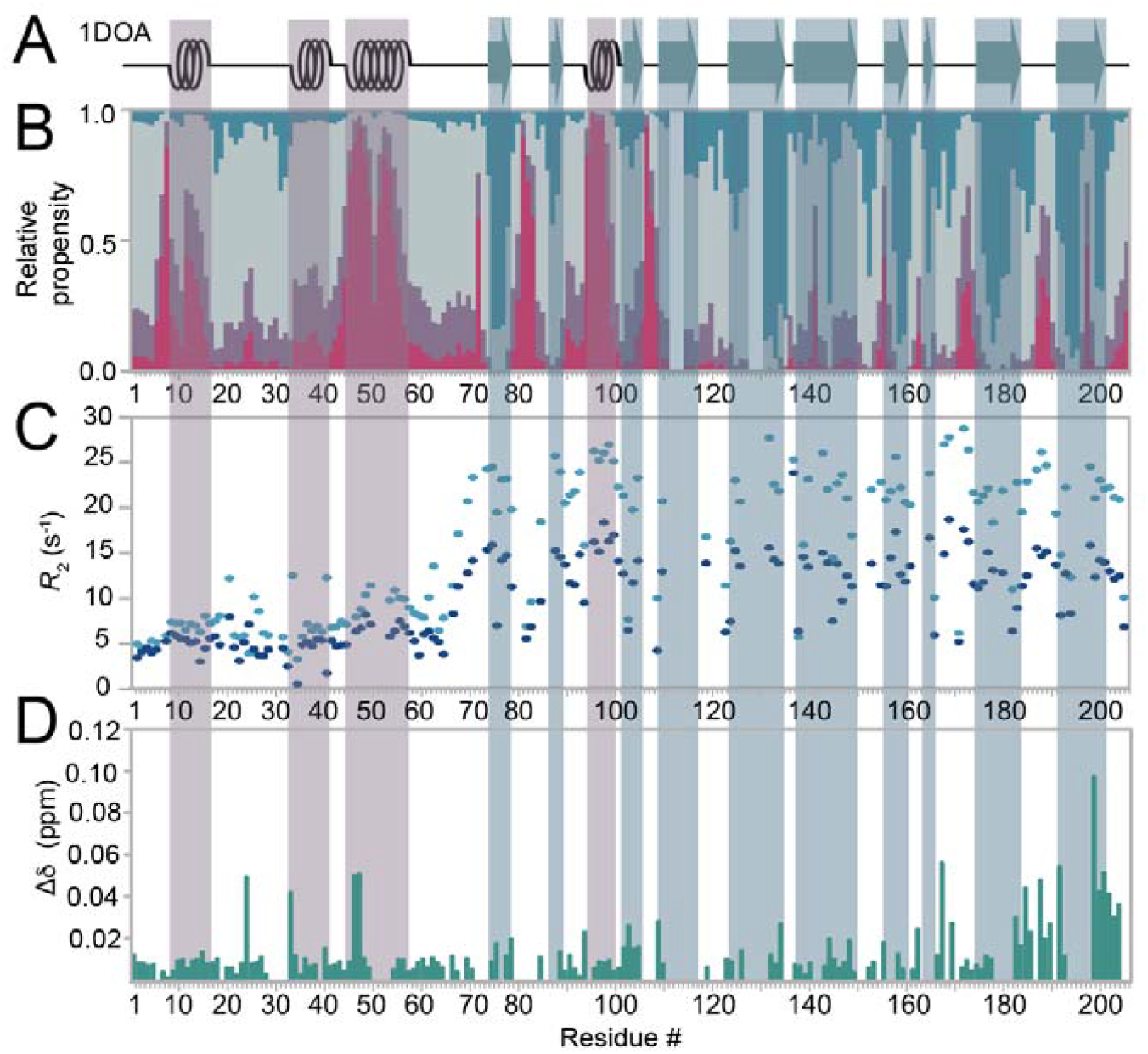
Structure and dynamics of the RhoGDI1 N-terminus in the GDI:Cdc42 complex. **A**) Secondary structure of the complex seen in 1DOA. **B**) Secondary-structural propensities of the complex derived from experimental chemical shifts (CheSPI). **C)** *R*_2_ rates of the complex (cyan), overlaid with those of the apo form (dark blue). D) Chemical-shift perturbations upon complex formation, calculated as (Δδ^²^(^1^ H)^N^ +Δδ^²^(^15^ N)/100)^0.5^

Even though the X-ray structure of the complex (PDB 1DOA) converges to a defined structure of the N-terminus, the electron density already reveals a certain degree of flexibility even under cryogenic conditions in the crystal (compare Fig. S6). C^α^ B-factors, reaching down to 34 within the remainder of the structure, bear values well above 100 up to residue 25 and between 59 and 66. Representing close-to-physiological conditions in solution, the NMR assessment at room temperature now reveals a highly mobile behavior of the N-terminus – even in the complex. The increase in *R*_2_ rates of representative residues in the inside of the core region agrees with what is expected for a complex of 46.6 kDa molecular weight at 25 °C (see Fig. 5C and also compare Fig. S7). In addition to the systematic overall increase of *R*_2_ rates of the N-terminal residues (from residue 7 onwards), a stronger increase is observed for those stretches with higher relative secondary-structural propensity (to well above 10 s^−1^ for the residues around residue 50). The largely retained dynamic behavior contradicts the stable folding onto the GTPase assumed in the bound state hitherto and rather demonstrates interactions with the client that retain a high degree of conformational freedom in the N-terminus. Adding to this picture of a rather loose association of the N-terminus and its preformed structural elements, only minor secondary-structural changes upon complex formation are apparent (Fig. 5B, compare with Fig. 3C). Apart from the moderate increase in most N-terminal *R*_2_ rates, significant local chemical-shift perturbations are observed (e. g., for residues V25, K33, E46, and S47, Fig. 5D) upon complex formation, matching with the expected N-terminal interactions between Cdc42 and RhoGDI1.^4^ In line with these observations, these sites tend to show a particular increase of the *R*_2_ rates upon complex formation (Fig. 5C), likely due to chemical-exchange contributions associated with these contacts. Together, these observations suggest a loose, plastic interaction of the GDI N-terminus with the binding partner and the remainder of the complex, with a remaining flexibility of most N-terminal residues, only few specific contacts, and the intrinsic secondary-structure distribution further maintained upon intermediate-timescale association/dissociation dynamics. Further characterization of the transient contacts between the GDI N-terminus and Cdc42 from the client side would be interesting. This is currently limited, however, by the incompatibility of the required geranylation of Cdc42 with either, isotope labeling or paramagnetic spin labeling.

Finally, we aimed to assess biochemically the importance of the internal structural stability of the GDI N-terminus for enhanced GTPase binding. On the basis of the above MD simulations and NMR data, we searched for a residue that is not part of the binding interface itself – and hence does not impact the binding affinity directly through intermolecular interactions with Cdc42 – but would only impact the tertiary structure of the temporary helix-loop-helix structural element within the GDI N-terminus. To identify our best candidate, we performed an alanine-scanning of the interface using the Robetta server^23^ (see Fig. S8). Fig. 6A visualizes the embedding of residue L56 in the helix-helix interactions via hydrophobic contact to I35. We chose decreasingly conservative mutations from Leu56 to Val, Ala, or Gly and read out the kinetic stability of the complex by a Förster resonance energy transfer (FRET)-based dissociation assay. In brief, the FRET signal of a dual-labelled complex (80 nM) with an intermolecular FRET pair (an Alexa647 acceptor on RhoGDI1 and a Cy3 donor on Cdc42, respectively) decreases upon mixing with excess (5 µM) unlabeled GDI, hence reporting on complex dissociation as a function of time. Indeed, we observed a strong and consistent increase in the dissociation rate constant of the complex upon perturbation of the inter-helical interaction with decreasing side chain length (Fig. 6B, C). These measurements, together with the association rate constant we independently determined for wildtype GDI (k_+GDI_ = 5 s^−1^ µM^−1^, Suppl. Fig. 9), allowed us to estimate the changes in complex affinity (Fig. 6C). We assumed that the GDI mutations cannot majorly impact the fast association rate constant for complex formation, which falls close to the diffusion limit and therefore cannot become much faster. Hence, we observed as previously^24^ that wildtype GDI bound its GTPase clients with extremely high affinity (K_D_ = 90pM) (Fig. 6C). Mutations of L56 lead to a progressive loss in affinity up to more than 400-fold. However, even the most weakly binding GDI variant (L56G) still possessed high affinity (K_D_ = 4 nM) for prenylated Cdc42.

**Fig. 6:**
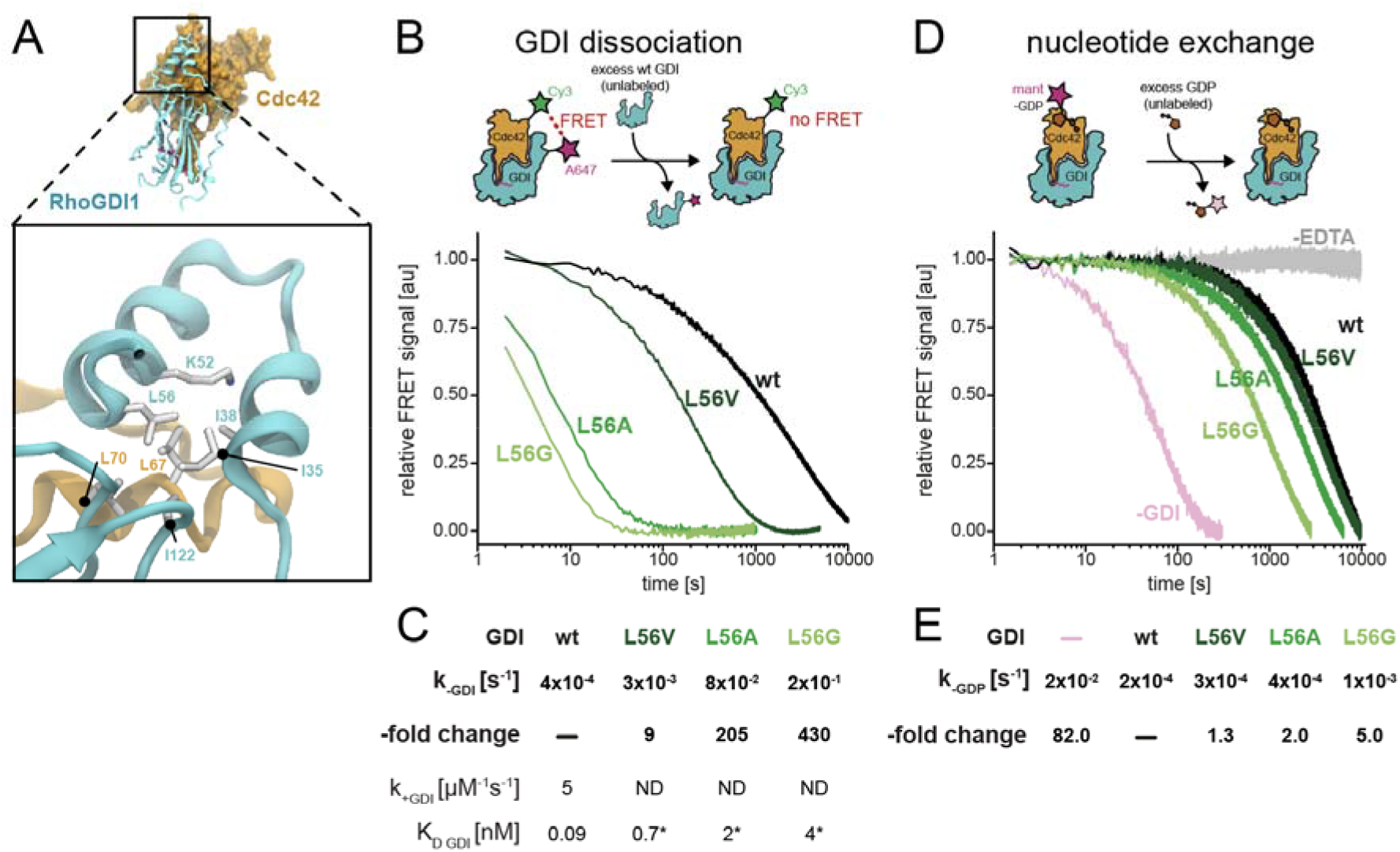
Assessment of the modulation of complex stability by destabilization of intramolecular interactions in the RhoGDI N-terminus via FRET. **A**) Environment of L56 within the overall structure (top) and locally within the C-terminal end of the helix-loop-helix motif (bottom). **B**) Top: Scheme of the FRET-based dissociation experiment. A 1:1 complex between Cy3-labeled, prenylated Cdc42 and Alexa647-labeled RhoGDI1 (wt or mutant) is mixed with excess wt RhoGDI1. The dissociation of the complex is followed over time by the loss in FRET between the two fluorophores. Bottom: Decrease in FRET signal as a function of time for wt RhoGDI (black) and mutants L56V, L56A, and L56G (dark to light green) as indicated. **C**) Summary of the dissociation rates (k_-GDI_) of GDI (either wt or mutants as indicated) from prenylated Cdc42 obtained from mono-exponential fits to the data shown in B and the relative rate enhancement compared to wildtype GDI. The corresponding association (k_+GDI_) rate was measured for wildtype GDI (see Fig. S. 9) and assumed to be the same or slower for the GDI variants. Equilibrium dissociation constants (K_D GDI_) were calculated by the rate of the measured dissociation rate constants and the measured or assumed association rate constant. **D**) Top: Scheme of the fluorescence intensity-based nucleotide exchange experiment. Prenylated Cdc42, loaded with Mant-GDP was mixed with excess (5 µM) RhoGDI1 (wt or mutant) to ensure continuous RhoGDI binding during the experiment. Nucleotide dissociation was initiated by the addition of excess unlabeled GDP (100µM) and followed over time by the loss in fluorescence intensity due to solvent exposure of the Mant-GDP upon unbinding from Cdc42 in complex with RhoGDI. Bottom: Decrease in Mant fluorescence as a function of time for either free prenylated Cdc42 (pink) or in complex with RhoGDI (either wildtype (black) or mutants L56V, L56A, and L56G (dark to light green)) as indicated. **E**) Summary of the nucleotide exchange rates (k_-GDP_) of Mant-GDP from prenylated Cdc42 either free or in complex with RhoGDI (either wt or mutants as indicated) obtained from mono-exponential fits to the data shown in D.

One established function of the GDI N-terminus is the inhibition of nucleotide exchange and hydrolysis in its bound GTPase.^4, 6, 7, 8^ Hence, we wondered how the intrinsic N-terminal dynamics and their enhancement by the destabilizing mutations in GDI affect the kinetics of GTPase nucleotide exchange. To this end, we measured EDTA-induced nucleotide dissociation using fluorescently-labeled GDP analogue Mant-GDP bound to prenylated Cdc42 and in the presence of excess unlabeled GDP in solution (Fig. 6D). We conducted these experiments in the presence or absence of saturating (5 µM) concentrations of RhoGDI (either wildtype of mutants) to generate the fully GDI-bound or GDI-free state during the experiment. We observed that binding of wildtype GDI strongly (83-fold) inhibited nucleotide exchange in Cdc42 as expected (Fig. 6D). However, the rate of nucleotide exchange (Fig. 6D, k _–GDP_ = 2×10^−4^ s^−1^) was similarly slow as GDI dissociation (k _-GDP_= 4×10^−4^ s^−1^), showing that these processes occur on a similar time scale for wildtype GDI. This means that the pronounced N-terminal dynamics we observe for GDI within the complex, are orders of magnitudes faster. (The N-terminus has a shorter correlation time than the protein core, a property on the ns timescale.) Steric blockage by the N-terminus does hence not seem to constitute the bottleneck for nucleotide exchange, which is an inherently slow process (k _-GDP_= 2×10^−2^ s^−1^ even in the absence of GDI). By contrast, we found that destabilizing the N-terminal structure through mutations of L56 resulted in a (up to 5-fold) acceleration of nucleotide dissociation in proportion to the severity of the perturbation (Fig. 6 D,E), in congruency with the decreased lifetime of the GTPase:GDI complex. Nucleotide exchange hence does not seem to be influenced by the N-terminus other than the differential lifetime of the protein:protein complex. (The discrepancy between the larger acceleration of the protein complex dissociation, with rates up to 2×10^−1^ s^−1^, compared to a smaller acceleration of nucleotide exchange, up to 1×10^−3^ s^−1^, results from the fact that acceleration of nucleotide exchange by mutation is convoluted with the exchange rate of free Cdc42 of maximally 2×10^−2^ s^−1^.) The known effect of nucleotide exchange being slowed down by the GDI – despite the lack of sufficiently long-lived direct interactions with the nucleotide binding site – seems to owe to overarching features of complex formation, like a modulation of GTPase internal dynamics (e. g., stalling of breathing motion) upon GDI binding, otherwise facilitating exchange in the free GTPase case. It cannot, however, derive from a steric blocking of exchange via the GDI N-terminus.

## Discussion

The behavior witnessed in the assessment of the RhoGDI structural properties pursued here differs from the long-standing assumption of a disorder-to-order transition upon complex formation with its GTPase client. Instead, in the full-length protein in its apo state, the N-terminus contains most of the helix-loop-helix interface as a stably preformed structural element just as it is later found in the complex. Conversely, a high degree of flexibility remains in the N-terminal residues even after complex formation. This is true even though expected chemical-shift perturbations and an overall increase of *R*_2_ rates unambiguously confirm the formation of the heterodimeric complex.

Intrinsically disordered regions have often been observed to adopt local and/or temporary secondary structure, with important effects on the binding properties, affinity and selectivity towards binding partners.^25, 26, 27, 28^ This potentially includes the possibility of fine-tuning the interaction as a function of external events, which can sensitively modulate the structural propensities and thus influence the occurrence of downstream signaling events. Likewise, the presence of transient and externally tunable structural features in the RhoGDI N-terminus point to a more complex mechanistic role compared to the previously assumed disorder-to-order transition and the simple steric inhibition of nucleotide exchange. Instead, the observed fine-tuned balance of order and disorder, affording sufficiently defined external interaction surfaces dependent on the differential presence of internal interaction surfaces, fraternizes a further strong increase in binding affinity via the terminus with the requirement of high spatial plasticity for the N-terminal interactions. As such, the dynamic properties of the N-terminus appears as a highly versatile addition assisting the RhoGDI core in tight GTPase binding, with an external docking device triggering the accessibility of the interaction partner, compare “fly-casting”^29, 30^. Thermodynamically, an even higher degree of rigidity would allow for tighter binding. Such rigidification of the terminus did, however, not emerge in evolution, presumably because the affinity is already very high for a bimolecular interaction. Instead, while aiding complex stabilization only modestly via its residual structural properties, the retained flexibility of the terminus seems to be desirable for other reasons. Overall flexibility despite tunable local pre-ordering might be particularly important for membrane extraction of GTPases by RhoGDI, likely a multi-step process^11^, in which the available intermolecular contacts for the N-terminus can vary along the process. The prenyl moiety needs to be leveraged out of the membrane while the final binding interface of the GTPase is still inaccessible. Only then insertion of the prenyl group into the core of RhoGDI establishes binding of the GTPase in the way witnessed in 1DOA. This process seems difficult to reconcile, with either a high-affinity yet fully rigid or a fully unstructured but low-affinity N-terminus. We propose that membrane extraction instead requires a predefined but loosely attached auxiliary structure (compare Fig. 7A, top row) responsive to the momentary situation. Context-specific structural modulation of an overall flexible N-terminus could also avoid unspecific binding of other membrane-bound small GTPases to RhoGDI, while still conserving to a maximum degree the structural definition required for high-affinity binding to intended clients. This type of selectivity might be crucial, considering the high similarity between distinct classes of the Ras superfamily. Specific transient-structural and dynamic requirements for the N-terminal sequence are in line with its high degree of sequence conservation, which has been noteworthy in the light that only a small fraction of the residues are thought to undergo direct interaction with the client.

**Fig. 7:**
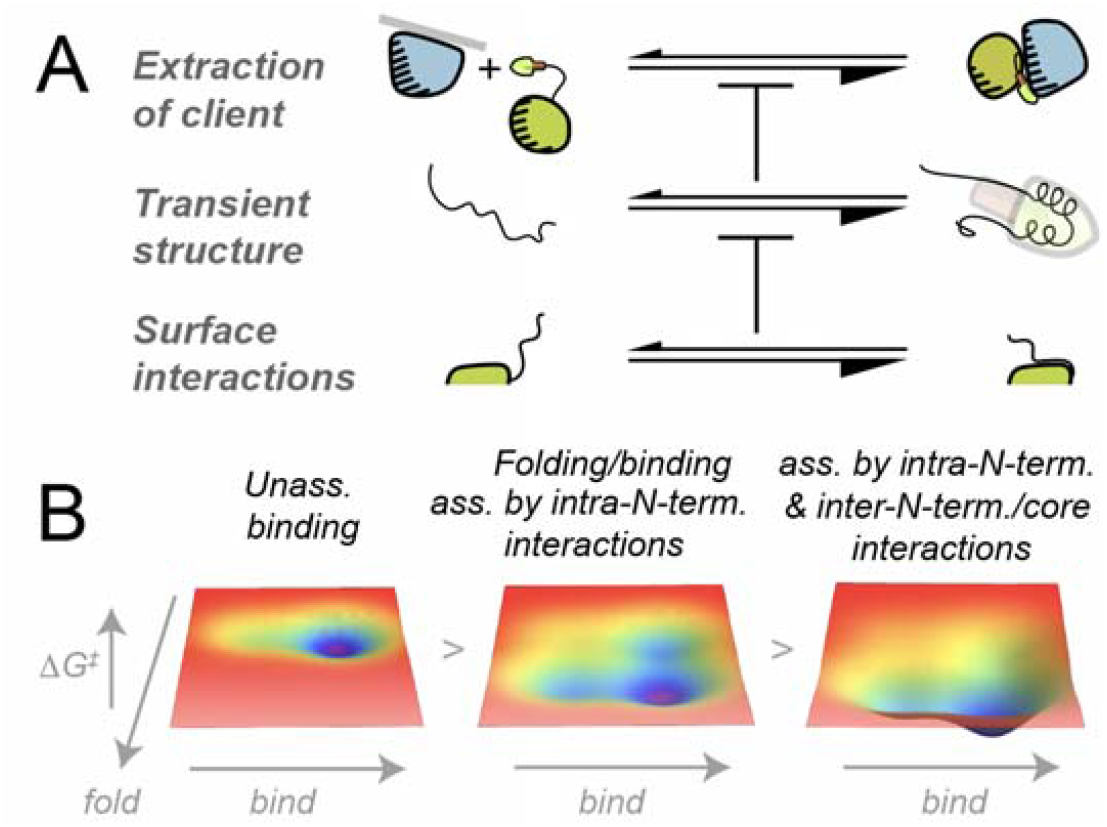
Thermodynamic interpretations of the N-terminal residual structural properties. **A**) In addition to a kinetic advantage of having a rather defined but still plastic *auxiliary structure* (represented as a tool flexibly attached by a string) for extraction/complex formation, a high tendency towards defined structural properties of the motif beneficial for its ability to capture the client is favorably modulated by transient interactions with the core (RhoGDI: green, client: blue). **B**) Comparative visualization of increasing facilitation of extraction, i. e., the potential-energy surfaces of simple binding of a disordered N-terminus without folding (left), coupled folding/binding in the presence of stabilizing intra-N-terminal interactions (middle), and coupled folding/binding in the favorable presence of cooperative interactions with nearby surfaces (right), further facilitating the folded and folded/bound states of the RhoGDI N-terminus while reducing entropic penalties of client association. Facilitated complex formation is assumed to concern not the final (sufficiently high) GTPase:GDI affinity, but critical transition states upon encounter with firmly membrane-tethered GTPase (ΔG^‡^).

Thermodynamically, the strong contribution to binding affinity by the N-terminus has been difficult to reconcile with a mechanistic model in which i) defined structural features of the motif are absent and ii) a large entropic penalty to the binding free energy would apply upon rigidification. The well-defined local intrinsic prefolding of the most important elements and the largely retained flexibility of the N-terminus revealed here show that neither of these drawbacks are actually present. The most peculiar biophysical aspect in addition to what has been described for intrinsically disordered regions in the context of coupled folding/binding events^31^ are the strongly differential (residual) structural properties of the N-terminal binding interface in the presence or absence of interactions with surfaces of the core (compare Fig. 4), pointing to a sensitive modulation of the folding energy landscape by transient interactions with seemingly innocent surfaces nearby (Fig. 7). The intramolecular surface interactions, the relative likelihood of which would differ upon membrane and client encounters, seem to suitably influence the residual N-terminal tertiary structural properties (i. e., towards a higher tendency of prefolding), which in turn facilitate the high-affinity binding of the client upon first contact, while maintaining plasticity. Similar effects have been described in the context of cooperative folding of nearby individual domains^32^ and the modulation on the free-energy surface by crowding agents and other temporarily interacting surfaces in a cellular environment^33^.

The data show that reconciliation of the seemingly contradictory picture of GDI-GTPase interactions requires the – experimentally verified – assumption of a fine-tuned co-existence of structure and disorder of the N-terminus in a context-specific equilibrium, modulated by both, interactions within the terminus as well as interactions to nearby surfaces. Bearing peculiar opportunities both, for nature and upon human interference to tweak affinity and selectivity, the large spectrum of different degrees of this co-existence, of importance for complex formation in a range of different areas of structural biology,^15, 34, 35, 36^ only begins to be understood.

## Conclusion

Here we used NMR spectroscopy in solution to capture the contradictory structural properties of the N-terminus of the Rho guanine nucleotide dissociation inhibitor in the absence and presence of its GTPase client. Opposed to a paradoxical disorder-to-order transition long assumed upon complex formation, the data demonstrate a context-specific propensity of the binding interface to preform those structural features later observed in the binding of the apo RhoGDI to its GTPase. Apart from suitable intra-terminal interactions constituting a (very modest) folding tendency of the N-terminal binding interface, pronounced structural features required for interface formation are effectively orchestrated by the transient interactions between terminus and core surface residues. Conversely, the overall degree of flexibility for this domain remains rather high even upon complexation. Together, the data point to a context-specific behavior and fine-tuned role of the N-terminus and its associated contact interfaces in the multi-step process of capturing and tethering the GDI clients opposed to forming an induced binding interface in the framework of coupled binding/folding or sterics-derived functionalities.

## Materials and Methods

Doubly (^13^C, ^15^N) or triply labeled (^2^H, ^13^C, ^15^N) bovine RhoGDI1 (full length or the isolated N-terminus (residues 1-59) was expressed in BL21(DE3) E. coli cells as a GST fusion construct and purified largely according to published protocols as described in more detail in the SI Appendix. Human Cdc42 was expressed as a His^6^-TEV fusion construct, purified similarly to GDI and *in-vitro* prenylated using geranylgeranyl transferase type 1 and geranylgeranyl diphosphate and labeled as previously described ^3^. (See details in the SI.) NMR spectra were recorded at 298 K on an 800 MHz Bruker Avance NEO spectrometer with a proton-optimized TCI cryo probe. Resonance assignments were derived from 2D ^15^N^−1^H HSQC, 3D HNCA, 3D HNCO, 3D HN(CA)CO and HNCACB/CBCANH for each sample as described previously,^37^ as well as via an additional 3D ^15^N NOESY-HSQC and a 3D HNHA for the isolated N-terminus, using CcpNmr v3^38^. NMR chemical shifts have been deposited into the BMRB under accession code 51835. ^15^N relaxation data were obtained by standard experiments as described in more detail in the SI. CheSPI^20^ analysis for quantification of residual secondary structure was run using the Web server (https://st-protein.chem.au.dk/chespi) based on experimental chemical shift information for ^1^H^N^, N, CO, Cα, and Cβ. For the fluorescence-based assays, prenylated Cdc42 was labeled at the N-terminus with Cy3-LPTEGG via Sortase-mediated peptide ligation. For dissociation assays, dual-labeled complexes of Alexa647-RhoGDI1 (wt or mutants) and prenylated Cy3-Cdc42 were formed *in situ* by mixing both proteins at final concentrations of 10 µM and incubating at RT for 10 minutes before being diluted to 80 nM monitored in a PTI QM-6 spectrofluorimeter using a 570 and 680 nm wavelength for donor and acceptor fluorescence, respectively. Then, 5 µM unlabeled RhoGDI1 (62-fold excess over Alexa647-RhoGDI1) was added and the relative change in FRET was monitored as a function of time. (See the SI for more details.) For association assays, prenylated Cy3-labeled Cdc42 (200 nM) was rapidly mixed with excess Alexa647-RhoGDI1 (790-2500 nM) in a stopped-flow device under constant donor excitation (520 nm) while monitoring the sensitized emission signal through a 670/40nm bandpass filter over time. (See the SI for more details.) For nucleotide exchange assays, Mant-GDP-bound, prenylated Cdc42 was diluted to 100 nM in the presence or absence of excess RhoGDI1 (5 µM, wt or mutants) and unlabeled GDP in a spectrofluorimeter. Then, 10 mM EDTA was added and the relative change in Mant fluorescence was monitored at 440 nm as a function of time. (See the SI for more details.) Molecular dynamics simulations were pursued in GROMACS 2022^39^ as structure-based model (SBM) simulations as a first access to interrogate GDI structural properties, starting from the structure of RhoGDI1 in complex with Cdc42 (PDBID: 1DOA^4^). (See further details in the SI.) A total of 20 simulations were performed, where 10 started from the fully folded state and 10 from a RhoGDI model with unfolded N-terminus, with a temperature range between 0.9 – 1.1 × the melting temperature. Under these conditions, the core of RhoGDI remains properly folded during the entire simulation. VMD^40^ was used to analyze the number of native contacts along the trajectories. A contact was considered to be formed if the two atoms were at a distance smaller than 1.2 × the one observed in the reference structure. Free-energy profiles were obtained using the weighted-histogram analysis method^41^.

## Acknowledgements

Funded by the Deutsche Forschungsgemeinschaft (DFG, German Research Foundation) individual grants 27112786, 325871075 to R.L. and 399893760 to P.B. and the Emmy Noether program to R.L. Funded by the Deutsche Forschungsgemeinschaft (DFG, German Research Foundation) under Germany’s Excellence Strategy -EXC 2033 – 390677874 – RESOLV, and EXC-114 – 24286268 – CiPS-M.

